# Direct acetylation of the conserved threonine of an immune regulator by bacterial effectors activates RPM1-dependent immunity in *Arabidopsis*

**DOI:** 10.1101/2021.02.05.426302

**Authors:** Sera Choi, Maxim Prokchorchik, Hyeonjung Lee, Ravi Gupta, Yoonyoung Lee, Buhyeon Cho, Min-Sung Kim, Sun Tae Kim, Kee Hoon Sohn

**Author notes:** These authors contributed equally: Sera Choi, Maxim Prokchorchik.

## Abstract

Plant pathogenic bacteria deliver effectors into plant cells to suppress immunity and promote pathogen survival (Buttner, 2016; Deslandes and Rivas, 2012); however, these effectors can be recognised by plant disease resistance (R) proteins to activate innate immunity (Jones and Dangl, 2006; Spoel and Dong, 2012). The bacterial acetyltransferase effectors HopZ5 and AvrBsT trigger immunity in *Arabidopsis thaliana* genotypes lacking *SUPPRESSOR OF AVRBST-ELICITED RESISTANCE 1* (*SOBER1*) (Choi et al., 2018; Jayaraman et al., 2017),. Using an *Arabidopsis* accession, Tscha-1, that naturally lacks functional *SOBER1* but is unable to recognise HopZ5, we demonstrate that *RESISTANCE TO P. SYRINGAE PV MACULICOLA 1* (*RPM1*) and *RPM1-INTERACTING PROTEIN 4* (*RIN4*) are indispensable for HopZ5- or AvrBsT-triggered immunity. Remarkably, T166 of RIN4, the phosphorylation of which is induced by AvrB and AvrRpm1, was directly acetylated by HopZ5 and AvrBsT. Furthermore, we demonstrate that acetylation of RIN4 T166 is required and sufficient for activation of HopZ5- or AvrBsT-triggered defence. Finally, we show that SOBER1 interferes with HopZ5- or AvrBsT-triggered immunity by deacetylating RIN4 T166. Our findings indicate that multiple pathogen effectors with distinct biochemical properties modify a single residue in a guardee protein and activate a plant NLR immune receptor. We have thus elucidated detailed molecular mechanisms underlying the activation and suppression of plant innate immunity triggered by bacterial acetyltransferases.

## Introduction

Many gram-negative bacterial pathogens deliver effector proteins into plant cells via a type III secretion system (T3S). Bacterial effectors modify host targets to interfere with plant innate immunity, resulting in successful colonization of host plants (Buttner, 2016; Deslandes and Rivas, 2012). In turn, plants have evolved intracellular disease resistance (R) proteins, mostly represented by nucleotide-binding (NB) and leucine-rich repeat (LRR) domain-containing (NLR) proteins, that recognise the corresponding effectors and activate immune responses (Jones and Dangl, 2006; Spoel and Dong, 2012). Some NLRs detect host target modifications caused by multiple avirulence effectors (Cesari et al., 2013; Grant et al., 1995; Narusaka et al., 2009). For example, the *Arabidopsis thaliana* NLR protein RPM1 (RESISTANCE TO PSEUDOMONAS SYRINGAE PV. MACULICOLA 1) recognises the *Pseudomonas syringae* effectors AvrRpm1 (ADP-ribosyl-transferase) and AvrB by sensing the phosphorylation status of RIN4 (RPM1-INTERACTING PROTEIN 4) during infection (Grant et al., 1995; Mackey et al., 2002; Redditt et al., 2019).

The *P. syringae* pv. *actinidiae* effector HopZ5 is an acetyltransferase that belongs to the YopJ effector family (Jayaraman et al., 2017; McCann et al., 2013). HopZ5 activity induces immune responses, including the hypersensitive response (HR) in the *Arabidopsis* Ct-1 accession but not in Col-0 due to the presence of a suppressor gene, *SUPPRESSOR OF AVRBST-ELICITED RESISTANCE 1* (*SOBER1*) (Choi et al., 2018). SOBER1 is a deacetylase that suppresses immune responses triggered by several acetyltransferase avirulence effectors, including HopZ5 and AvrBsT (Burger et al., 2017; Choi et al., 2018; Cunnac et al., 2007). Among several AvrBsT-interacting proteins identified in pepper and Arabidopsis, SOBER1 deacetylates the AvrBsT-mediated acetylation of its target protein ACETYLATED INTERACTING PROTEIN 1 (ACIP1), which is partially required for AvrBsT-triggered immunity (Burger et al., 2017; Cheong et al., 2014; Han and Hwang, 2017). However, the detailed molecular mechanisms by which SOBER1 suppresses AvrBsT- or HopZ5-triggered immunity are not fully understood mainly because the detailed molecular mechanisms for AvrBsT- or HopZ5-triggered immunity are not fully understoood.

Here, we show that both HopZ5 and AvrBsT directly acetylate the conserved RIN4 threonine residue T166 during bacterial infection. RIN4 T166 acetylation is necessary and sufficient forRPM1-dependent immunity activation. Importantly, we also show that SOBER1 deacetylation of RIN4 T166 acetylation results in suppression of HopZ5- or AvrBsT-triggered immunity. Taken together, we reveal the detailed molecular mechanisms underlying the activation and suppression of bacterial acetyltransferase effector-triggered immunity.

## Results and Discussion

### HopZ5 or AvrBsT-triggered immunity is *RPM1*-and *RIN4*-dependent

Bacterial T3S-delivered HopZ5 triggers immune responses in the *Arabidopsis* genotypes lacking a functional *SOBER1*, such as Ct-1 (naturally lacking *SOBER1*) and the Col-0 *sober1-3* mutant, indicating the presence of a corresponding *R* gene(s) in both genotypes (Choi et al., 2018). To identify an *Arabidopsis* accession that lacks a functional *SOBER1* but which fails to show HopZ5-triggered immune responses, we tested the HopZ5 responsiveness of several *Arabidopsis* accessions (Es-0, Kyoto, and Tscha-1) predicted to carry a nonsense mutation in *SOBER1* (signal.salk.edu/atg1001/3.0/gebrowser.php). Among these accessions, only Tscha-1 failed to show a HR in response to HopZ5 delivered from *P. fluorescens* Pf0-1 carrying a functional T3S (hereafter, Pf0-1) (Thomas et al., 2009) (Figure 1A). Consistent with the HR result, the *in planta* growth of *P. syringae* pv. *tomato* (*Pto*) DC3000 carrying *hopZ5* was comparable to a catalytically inactive variant, *hopZ5*^*C218A*^, or an empty vector (EV) in Tscha-1 (Figure 1B). These results indicate that Tscha-1 may lack the corresponding *R* gene for HopZ5.

**Figure 1.**
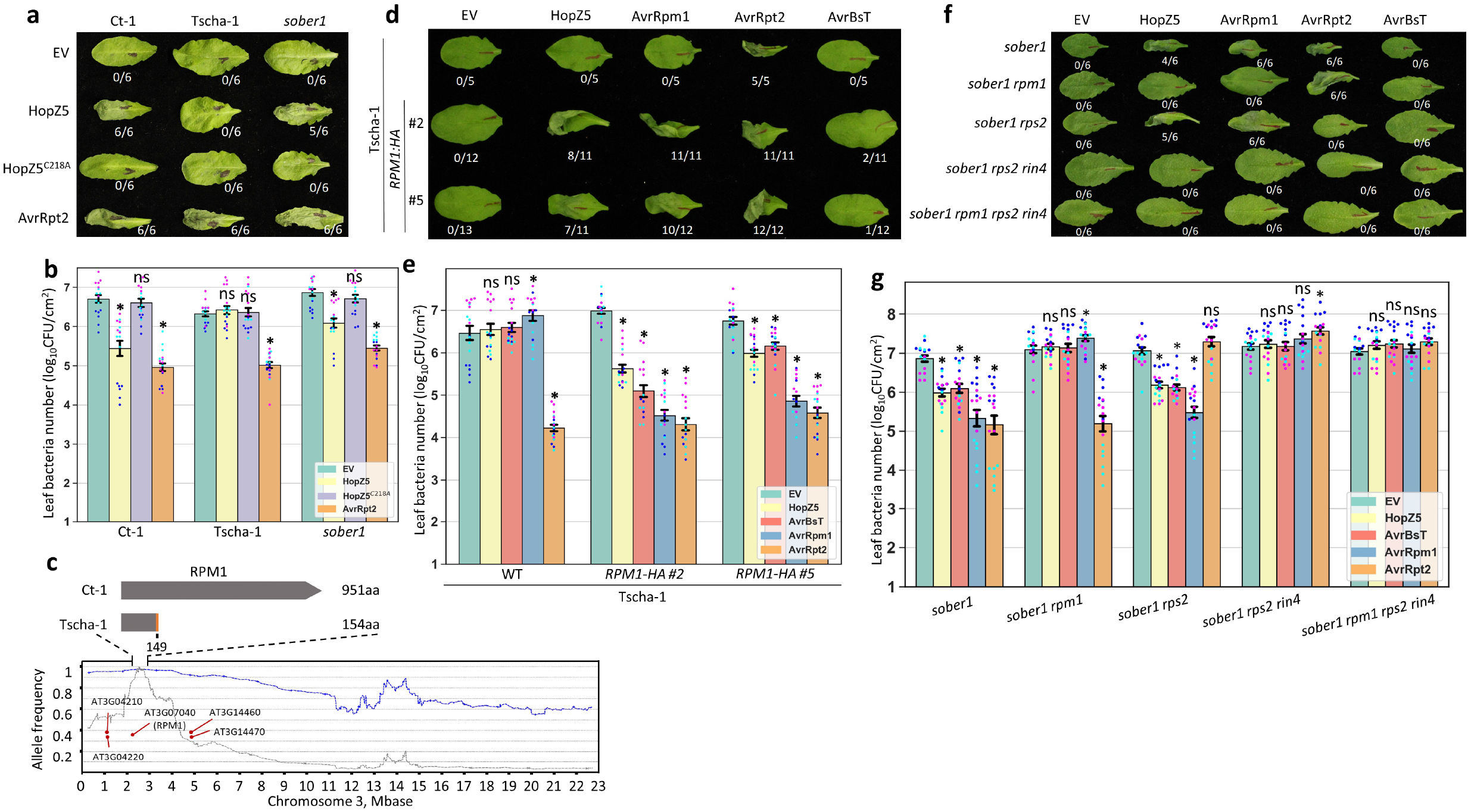
Bacterial acetyltransferase effectors HopZ5 and AvrBsT trigger RPM1- and RIN4-dependent immunity in *Arabidopsis*. **(A)** HopZ5 triggers the hypersensitive response (HR) in *Arabidopsis* Ct-1 and *sober1*, but not in Tscha-1. HR symptoms developed after infiltrating with *Pseudomonas fluorescens* Pf0-1 (T3S) carrying EV, *hopZ5, hopZ5*^*C218A*^, or *avrRpt2* were scored and photographed at 1 day post infiltration (dpi). (**B)** HopZ5 restricts *Pto* DC3000 growth in *Arabidopsis* Ct-1 and *sober1* but not in Tscha-1. The growth of *Pto* DC3000 carrying EV, *hopZ5, hopZ5*^*C218A*^, and *avrRpt2* was quantified at 4 dpi. The data shown are the mean ± s.e.m. (**C)** Identification of the *RPM1* locus conferring HopZ5 recognition. The genomic DNA pool of *Arabidopsis* Ct-1 × Tscha-1 F_2_ plants which did not show HopZ5-triggered HR was sequenced and the SNP allele frequency (AF) was calculated. A window-based AF graph is shown as blue dashed line. The grey dashed graph represents the window-based boost AF values (window-size: 600,000, window step: 15,000, marker min score: 20, marker min coverage: 30). Gene ID numbers (from www.arabidopsis.org) indicate predicted *NLR* genes near the mapped interval. The *RPM1* gene in Tscha-1 encodes a severely truncated (154 amino acids) RPM1 protein with a frameshift from T149 (indicated in orange). (**D)** HopZ5 triggers the HR in transgenic Tscha-1 lines expressing *RPM1*_*Col-0*_. HR symptoms developed after infiltrating Pf0-1 carrying EV, *hopZ5, avrRpm1, avrRpt2*, or *avrBsT* were scored at 1 dpi. HopZ5- or AvrBsT-triggered HR was observed in two independent *RPM1-HA*-expressing transgenic Tscha-1 lines (#2 and #5). (**E)** The transgenic expression of *RPM1* confers the HopZ5- or AvrBsT-triggered restriction of *Pto* DC3000 growth. The growth of *Pto* DC3000 carrying EV, *hopZ5, avrRpm1, avrRpt2*, or *avrBsT* was quantified at 4 dpi. The data shown are the mean ± s.e.m. **F**, HopZ5-triggered HR requires RPM1 and RIN4. The HR was tested in *sober1, sober1 rpm1, sober1 rps2, sober1 rps2 rin4*, and *sober1 rpm1 rps2 rin4*, as described for **D**. (**G)** The HopZ5- and AvrBsT-triggered restriction of *Pto* DC3000 growth is dependent on RPM1 and RIN4. *Pto* DC3000 growth restriction was tested in *sober1, sober1 rpm1, sober1 rps2, sober1 rps2 rin4*, and *sober1 rpm1 rps2 rin4*, as described for **E** The data shown are the mean ± s.e.m. In **A, D**, and **F**, the numbers indicate the number of leaves showing the HR/total infiltrated leaves. In **B, E**, and **G**, the coloured dots represent different biological replicates (*n* = 6). Asterisks and ns mean Student t-test *p*-value < 0.05 or > 0.05 compared to EV, respectively. All experiments in this figure were conducted at least three times with similar results.

Next, we generated a Ct-1 × Tscha-1 F_2_ mapping population to identify the HopZ5-recognising *R* gene. In this population, the HopZ5-triggered HR phenotype segregated at a 3:1 (HR:no HR) ratio, suggesting that a single dominant *R* locus confers HopZ5-triggered immunity (Table S1). Wild-type parental accessions, Ct-1 and Tscha-1, and the pool of 35 F_2_ plants that did not show HopZ5-triggered HR were subjected to next-generation sequencing. A single nucleotide polymorphism (SNP) allele frequency analysis in the pool of F_2_ plants that failed to show HopZ5-triggered HR indicated a dramatic enrichment of Tscha-1-specific SNPs on the upper arm of chromosome 3 in the ∼600-kbp interval between 2.2 and 2.8 Mbp (Figure 1C). Among the five *NLR* genes in this region (Figure 1C), only *RPM1* carried a nonsense mutation (single nucleotide deletion) resulting in a premature stop codon in Tscha-1 but not in Col-0 or Ct-1 (Figure 1C) (Grant et al., 1995). To test whether RPM1 confers HopZ5 recognition, two independent transgenic Tscha-1 lines carrying *RPM1*^*Col-0*^, whose expression is driven by its native promoter, were developed and tested for HopZ5-triggered immunity (Figure S1). These transgenic lines showed a strong HR in response to Pf0-1-delivered HopZ5, but not to AvrBsT; however, in a bacterial growth assay, *Pto* DC3000 carrying *hopZ5* or *avrBsT* showed a RPM1-dependent reduction in growth, demonstrating that RPM1 confers the recognition of both avirulence effectors (Figure 1E). Consistent with our results, a highly overlapping recognition specificity between HopZ5 and AvrBsT was reported previously in *Arabidopsis*, with the exception of a differing HR phenotype in the *sober1* mutant (Choi et al., 2018; Cunnac et al., 2007).

Next, we set out an experiment to investigate whether HopZ5- or AvrBsT-triggered immunity requires RIN4, since it is required for RPM1 function in recognition of the *P. syringae* avirulence effectors AvrRpt2, AvrRpm1, and AvrB. To test the RIN4 requirement for HopZ5- or AvrBsT-triggered immunity, *sober1 rpm1, sober1 rps2, sober1 rps2 rin4*, and *sober1 rpm1 rps2 rin4* mutants were developed (see Methods for details). Since the *rin4* single mutant is not viable due to RPS2-mediated autoimmunity (Belkhadir et al., 2004; Mackey et al., 2003), *sober1 rps2 rin4* was developed instead of *sober1 rin4*. In the HR and *Pto* DC3000 growth restriction assays, AvrRpm1 or AvrRpt2 triggered RIN4-/RPM1- or RIN4-/RPS2-dependent immune responses, respectively, as previously reported (Bisgrove et al., 1994; Kunkel et al., 1993; Mackey et al., 2002). Consistent with our Tscha-1 (*RPM1:HA*) results, Pf0-1 expressing *HopZ5* but not *AvrBsT* triggered the HR in a RPM1- and RIN4-dependent manner (Figure 1F). Moreover, the growth of *Pto* DC3000 carrying *hopZ5* or *avrBsT* was comparable with those carrying the EV in the *sober1 rpm1* and *sober1 rps2 rin4* mutants, indicating that *RPM1* and *RIN4* are required for HopZ5- or AvrBsT-triggered immunity in *Arabidopsis* (Figure 1G). Taken together, these results suggest that the two closest YopJ family bacterial acetyltransferases, HopZ5 and AvrBsT, may target RIN4, resulting in an activation of RPM1-dependent immunity.

### HopZ5 and AvrBsT directly acetylate a conserved threonine residue of RIN4 and activate RPM1

Based on these results, we hypothesized that RIN4 could be a direct substrate of HopZ5 and AvrBsT acetyltransferases. To test this hypothesis, we first tested the *in planta* protein– protein association between RIN4 and either HopZ5 or AvrBsT using an *in planta* co-immunoprecipitation (Co-IP) assay. AvrRpm1-GFP, HopZ5-YFP, and AvrBsT-YFP, as well as the catalytically inactive variants HopZ5^C218A^-YFP and AvrBsT^H154A^-YFP, associated with FLAG-RIN4 when transiently expressed in *Nicotiana benthamiana* (Figure 2A). The catalytic activity of AvrB is unknown, but AvrRpm1 was recently reported to be an ADP-ribosyl-transferase (Desveaux et al., 2007; Redditt et al., 2019). Both effectors eventually induce the phosphorylation of RIN4 T166, which is fully or partially required for the RPM1-dependent recognition of AvrB or AvrRpm1, respectively (Chung et al., 2011; Desveaux et al., 2007; Liu et al., 2011). As HopZ5 and AvrBsT possess acetyltransferase activity (Cheong et al., 2014; Jayaraman et al., 2017), we set out an experiment to identify the acetylated amino acid residue(s) of RIN4. The FLAG-RIN4 recombinant protein was co-incubated with HopZ5-FLAG or HopZ5^C218A^*-*FLAG *in vitro* and subjected to a liquid chromatography (LC)–mass spectrometry (MS)/MS analysis, revealing that RIN4 T166 was the only acetylated residue that could be identified when incubated with wild-type HopZ5 but not HopZ5^C218A^ (Figure 2B and S2). For the further validation of T166 acetylation by HopZ5, a polyclonal antibody (α-AcRIN4) was raised against the acetylated T166 RIN4 fragment peptide (CGADGYTacHIFN) and used for the detection of T166-acetylated RIN4. To test whether HopZ5 or AvrBsT directly acetylates RIN4 T166, *in vitro–*purified FLAG-RIN4 was incubated with a wild-type or catalytically inactive variant of HopZ5 or AvrBsT. The wild-type HopZ5 and AvrBsT, but not HopZ5^C218A^ and AvrBsT^H154A^, acetylated RIN4 T166 in an immunoblot analysis using the α-AcRIN4 antibody (Figure 2C). Next, to determine whether RIN4 T166 is acetylated by HopZ5 and AvrBsT in plant cells, we transiently expressed MYC-RIN4, HopZ5-FLAG, and AvrBsT-FLAG in *N. benthamiana* leaves and performed a Co-IP assay. Consistent with our *in vitro* results, RIN4 T166 was acetylated when co-expressed with the wild-type but not catalytically inactive variant of HopZ5 or AvrBsT in *N. benthamiana* (Figure 2D). To test whether RIN4 is acetylated during bacterial infection, we infected transgenic *Arabidopsis* plants (*sober1*_*Ct-1*_ *rps2 rin4*) expressing FLAG-RIN4 with Pf0-1 carrying EV, *hopZ5*, or *avrB*. Importantly, the bacterial T3S delivery of HopZ5 but not AvrB resulted in the acetylation of RIN4 at 6 and 12 hours post infection (hpi) (Figure 2E). Together, our results demonstrate that HopZ5 and AvrBsT directly acetylate RIN4 T166 during bacterial pathogenesis.

**Figure 2.**
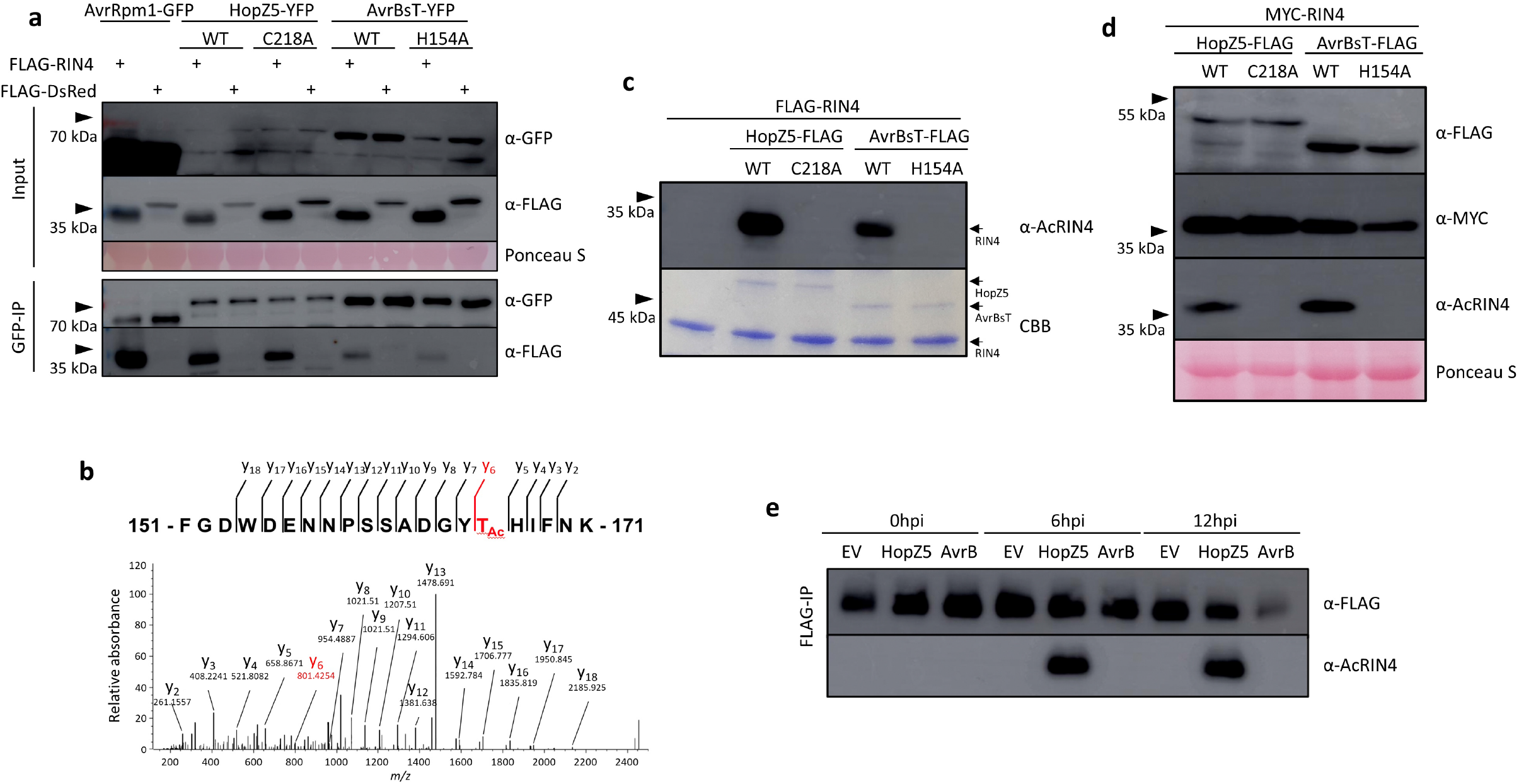
HopZ5 and AvrBsT interact with and acetylate RIN4. **(A)** HopZ5 and AvrBsT interact with RIN4 regardless of their catalytic activity. AvrRpm1-GFP, HopZ5 (wild-type (WT) or C218A mutant)-YFP, and AvrBsT (WT or H154A)-YFP proteins were co-expressed with FLAG-RIN4 or FLAG-DsRed in *Nicotiana benthamiana*. Total protein extracts and α-GFP-immunoprecipitated proteins were immunoblotted. Ponceau S staining of the rubisco protein band is shown as a loading control. **(B)** MS/MS spectra of the acetylated RIN4 fragment peptide (FGDWDENNPSSADGYTHIFNK) after incubation with HopZ5-FLAG protein. The y ions are marked in the spectrum and illustrated (m/z = 1229.02). (**C)** HopZ5 and AvrBsT acetylate T166 of RIN4 *in vitro*. FLAG-RIN4 was incubated with HopZ5 (WT or C218A)-FLAG or AvrBsT (WT or H154A)-FLAG and immunoblotted with an α-AcRIN4 antibody. The same amount of protein used in the immunoblot was loaded for Coomassie Brilliant Blue gel staining (CBB) in a separate gel. **(D)** HopZ5 and AvrBsT acetylate T166 RIN4 in *N. benthamiana*. HopZ5 (WT or C218A)-FLAG or AvrBsT (WT or H154A)-FLAG were co-expressed with MYC-RIN4 in *N. benthamiana*. The total protein extract was immunoblotted. Ponceau S staining of the rubisco protein band is a loading control. **(E)** HopZ5 acetylates T166 RIN4 in *Arabidopsis*. Leaf samples were collected from *sober1*_*Ct-1*_ *rps2 rin4* (*35S:FLAG:RIN4*) transgenic plants infiltrated with *P. fluorescens* Pf0-1 carrying EV, *hopZ5*, and *avrB* at each timepoint. Protein samples were α-FLAG-immunoprecipitated for immunoblot analysis. All experiments except **B** were conducted at least three times with similar results; **B** was conducted twice.

### RIN4 T166 acetylation is necessary and sufficient to activate RPM1-dependent immunity

We next investigated whether RIN4 T166 acetylation by HopZ5 or AvrBsT is required for the activation of RPM1-dependent immunity. Transgenic *sober1*_*Ct-1*_ *rps2 rin4* lines expressing the *FLAG:RIN4*^*T166A*^ mutant were generated to test the importance of T166 acetylation in RPM1 activation (Figure S3). Consistent with the previous report that RIN4 T166 phosphorylation is fully or partially required for AvrB-or AvrRpm1-triggered immunity (Chung et al., 2011), respectively, our transgenic lines expressing *FLAG:RIN4*^*T166A*^ developed a HR in response to Pf0-1-delivered AvrRpm1 but not AvrB (Figure 3A). Importantly, HopZ5 or AvrBsT triggered a robust HR in the transgenic *sober1*_*Ct-1*_ *rps2 rin4* plants expressing wild-type *FLAG:RIN4* but not *FLAG:RIN4*^*T166A*^ (Figure 3A). To test whether the RIN4^T166A^ mutation causes the loss of HopZ5- or AvrBsT-triggered restriction of virulent bacterial growth, transgenic *sober1*_*Ct-1*_ *rps2 rin4* plants expressing wild-type *RIN4* or *RIN4*^*T166A*^ were infected with *Pto* DC3000 carrying *hopZ5* or *avrBsT* (Figure 3B). Consistent with the HR assay results (Figure 3A), HopZ5- or AvrBsT-triggered *Pto* DC3000 growth restriction was abolished in *sober1*_*Ct-1*_ *rps2 rin4* (*FLAG:RIN4*^*T166A*^) (Figure 3B). RIN4 T166 acetylation is therefore indispensable for HopZ5- or AvrBsT-triggered RPM1-dependent immunity. Previously, the phosphorylation mimic mutant RIN4^T166D^ was reported to be sufficient to activate RPM1 in the absence of effectors (Chung et al., 2011; Liu et al., 2011; Xu et al., 2017); thus, we set out to investigate whether RIN4 T166 acetylation is sufficient to activate RPM1 by using a RIN4 variant that mimics T166 acetylation. Recently, it was shown that isoleucine can functionally mimic acetylated serine and possibly threonine (Bastedo et al., 2019). Based on this, we first tried to generate the transgenic *Arabidopsis* lines expressing *RIN4*^*T166I*^ or *RIN4*^*T166D*^, but failed to recover T_2_ generation seeds due to lethality (Figure S4) (Belkhadir et al., 2004; Mackey et al., 2003). Thus, we used the Tobacco rattle virus (TRV)-mediated transient protein overexpression system to express various mutant RIN4 proteins in *Arabidopsis* (see Methods for details) (Wroblewski et al., 2018). Due to the low efficiency of *Agrobacterium-*mediated transient expression in *Arabidopsis, N. benthamiana* leaf cells were infected with TRV derivatives carrying wild-type *RIN4, RIN4*^*T166A*^, *RIN4*^*T166I*^, or *RIN4*^*T166D*^ using this technique. Subsequently, the infected *N. benthamiana* sap was collected and used to infect *Arabidopsis*. In agreement with the previous results, the TRV-mediated expression of RIN4^T166I^ or RIN4^T166D^, but not wild-type RIN4 or RIN4^T166A^, resulted in RPM1-dependent cell death (Figure 3C). The protein expression levels of the RIN4 variants in infected *Arabidopsis* leaf cells were comparable, indicating that the lack of cell death was not due to the instability of the RIN4 protein (Figure 3D); RIN4^T166I^ is sufficient to activate RPM1-dependent defence. Remarkably, RIN4^T166I^-mediated cell death was not inhibited by the presence of SOBER1 in the *rps2 rin4* mutant, suggesting that SOBER1 cannot interfere with RPM1 activation when RIN4 carries an irreversible acetylation mimic (Figure 3C). Based on these results, we further hypothesized that SOBER1 suppresses HopZ5- or AvrBsT-triggered RPM1 activation by the deacetylation of a RIN4 T166 residue.

**Figure 3.**
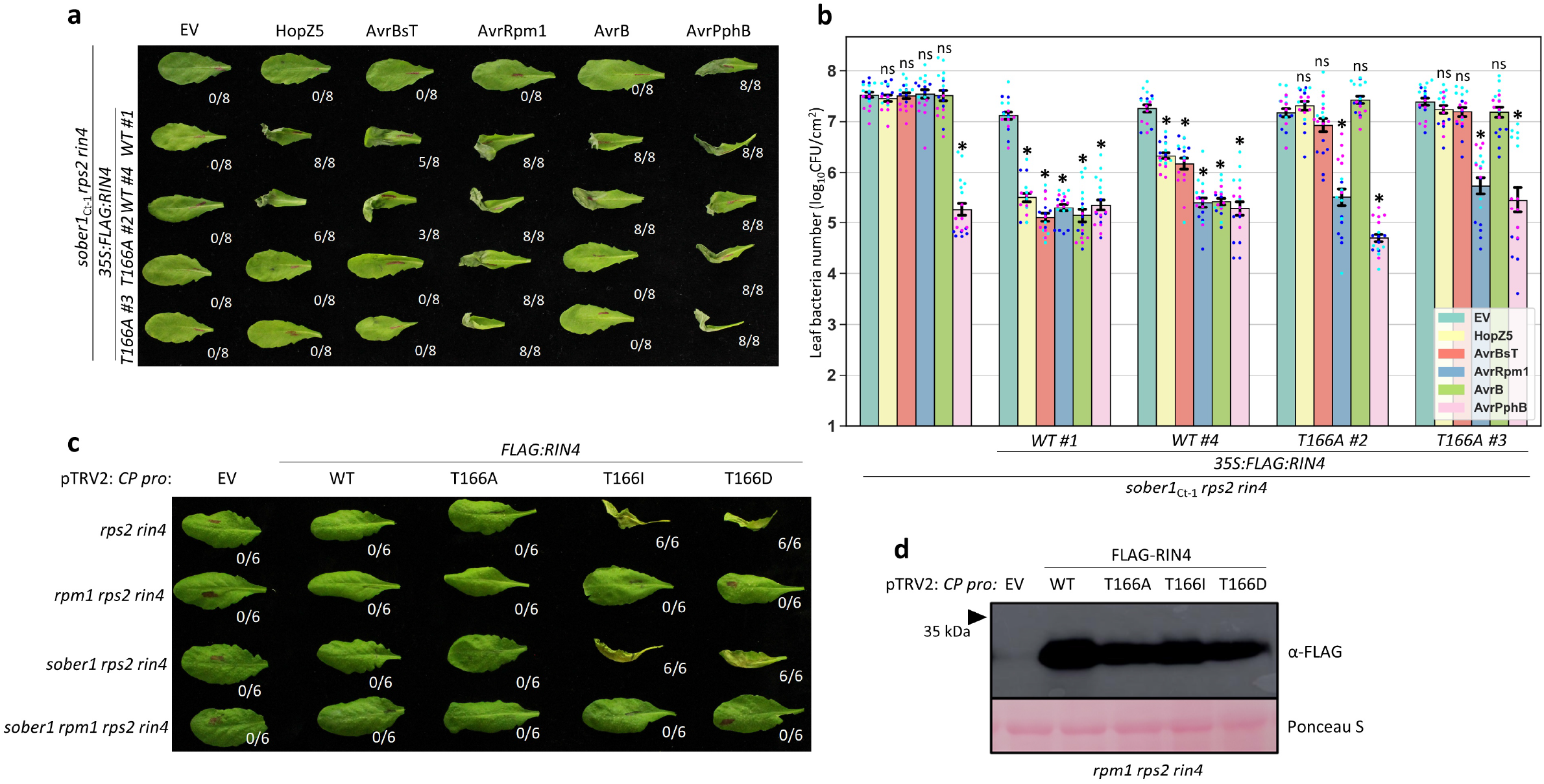
RIN4 T166 acetylation is indispensable and sufficient for HopZ5- or AvrBsT-triggered immunity. **(A)** RIN4 T166 is required for the HopZ5- or AvrBsT-triggered HR. HR symptoms developed after infiltrating Pf0-1 carrying EV, *hopZ5, avrBsT, avrRpm1, avrB*, or *avrPphB* were scored at 1 day post infiltration (dpi) in *sober1*_*Ct-1*_ *rin4 rps2* and two independent T_2_ transgenic lines of *sober1*_*Ct-1*_ *rin4 rps2* expressing *FLAG:RIN4* (WT or T166A). (**B)** RIN4 T166 is required for HopZ5- or AvrBsT-triggered restriction of *Pto* DC3000 growth. The growth of *Pto* DC3000 carrying EV, *hopZ5, avrBsT, avrRpm1, avrB*, or *avrPphB* was quantified at 4 dpi in the indicated plant genotypes. Coloured dots represent different biological replicates (*n* = 6). Asterisks and ns indicate Student’s t-test *p*-value < 0.05 or > 0.05 compared with EV, respectively. The data are shown as the mean±s.e.m. (**C)** TRV-mediated expression of RIN4 T166 acetylation mimicking the RIN4^T166I^ variant induces RPM1-dependent cell death in *Arabidopsis. Agrobacterium* harbouring pTRV1 and pTRV2:*CPpro:FLAG:RIN4* (WT, T166A, T166I, and T166D) variants-infected *N. benthamiana* sap was rub-inoculated with carborundum into *rps2 rin4, rpm1 rps2 rin4, sober1 rps2 rin4*, and *sober1 rpm1 rps2 rin4*. Cell death symptoms were scored at 7 dpi. (**D)** TRV-mediated protein expression of RIN4 variants in *Arabidopsis*. Leaf samples were taken at 2 dpi in *rpm1 rps2 rin4* as described in **C**, and total protein extracts were subjected to immunoblot analysis. Ponceau S staining of the rubisco protein band is shown as a loading control. In **A** and **C**, the numbers indicate the numbers of leaves showing cell death/total infiltrated leaves. All experiments were conducted at least three times with similar results.

### SOBER1 deacetylates RIN4 T166 acetylation

To test whether RIN4 is a substrate of the SOBER1 deacetylase, we first tested the association between transiently expressed SOBER1 and RIN4 in *N. benthamiana* using Co-IP. In agreement with our hypothesis, the wild-type SOBER1-FLAG and the catalytically inactive variant H192A (Choi et al., 2018; Cunnac et al., 2007) interacted with MYC-RIN4 (Figure 4A). Next, we used proteins purified from *Escherichia coli* to test the SOBER1-directed deacetylation of RIN4 T166 acetylation (Figure 4B). When tested with the anti-AcRIN4 antibody, the incubation of wild-type SOBER1-FLAG but not the H192A variant with FLAG-RIN4 and HopZ5-FLAG resulted in the loss of RIN4 T166 acetylation (Figure 4B), demonstrating that RIN4 is a direct enzymatic target of SOBER1. In addition, SOBER1-FLAG but not SOBER1^H192A^-FLAG deacetylated HopZ5- or AvrBsT-directed MYC–RIN4 T166 acetylation when transiently co-expressed in *N. benthamiana* (Figure 4C). These results strongly indicate that SOBER1 suppresses HopZ5- and AvrBsT-triggered immunity by deacetylating RIN4 T166 during bacterial pathogenesis.

**Figure 4.**
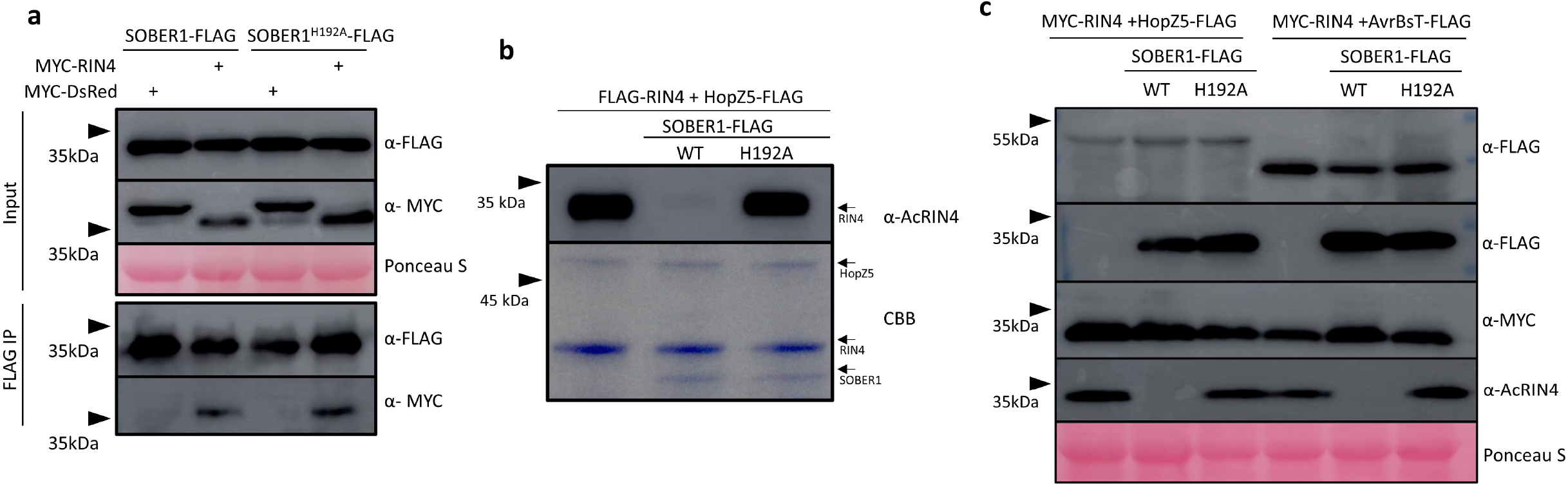
SOBER1 deacetylates HopZ5-directed RIN4 T166 acetylation. **(A)** SOBER1 interacts with RIN4 regardless of catalytic activity. SOBER1-FLAG or SOBER1^H192A^-FLAG proteins were co-expressed with MYC-RIN4 or MYC-DsRed in *Nicotiana Benthamiana* leaves using *Agrobacterium*-mediated transient transformation. The total protein extracts and α-FLAG antibody–immunoprecipitated proteins were analysed using immunoblotting. Ponceau S staining of the rubiscoprotein band is shown as a loading control. **(B)** SOBER1 deacetylates RIN4 T166 *in vitro*. FLAG-RIN4 was acetylated *in vitro* by HopZ5-FLAG in the presence of 50 μM acetyl-CoA and 100 nm IP_6_. SOBER1-FLAG or SOBER1^H192A^-FLAG was subsequently added before subjecting the sample to immunoblot analysis. The acetylation of RIN4 T166 was detected using the α-AcRIN4 antibody. The same amount of protein used in the immunoblot analysis was loaded for Coomassie Brilliant Blue (CBB) in a separate gel. (**C)** SOBER1 deacetylates RIN4 T166 when expressed in *Nicotiana benthamiana*. SOBER1-FLAG or SOBER1^H192A^-FLAG protein was co-expressed with MYC-RIN4 and HopZ5-FLAG or AvrBsT-FLAG in *N. benthamiana*. Total protein extracts were used for the immunoblot analysis using the indicated antibodies. Ponceau S staining of the rubisco protein band is shown as a loading control. All the experiments were conducted at least three times with similar results.

Plants must maximise the capability of their limited NLR protein inventory to recognise effectors with various biochemical activities (Toruno et al., 2016). One of the strategies to achieve this is to utilise guardee proteins (van der Hoorn and Kamoun, 2008). RIN4 is guarded by several NLRs and targeted by effectors with different biochemical activities (Chisholm et al., 2005; Chung et al., 2011; Mackey et al., 2003; Mackey et al., 2002; Redditt et al., 2019). Different modifications of RIN4 could be sensed by different NLRs (Axtell and Staskawicz, 2003; Chung et al., 2011); however, different biochemical modifications of the same amino acid residue of RIN4 or other guardee proteins that are sufficient to activate guarding NLRs have not been reported. Our results reveal a novel mechanism of RPM1 activation in which two bacterial effectors directly acetylate a conserved threonine residue of RIN4, whose phosphorylation is induced by AvrB and AvrRpm1. Thus, RIN4 T166 can be targeted by multiple effectors, enabling RIN4 to be an efficient guardee protein. In addition, the mechanistic details revealed in our study explain the specificity of SOBER1 function in suppression of effector-triggered immunity.

## Methods

### Plant materials and growth conditions

*Arabidopsis thaliana* Ct-1 and Tscha-1 were obtained from the Nottingham Arabidopsis Stock Centre (United Kingdom). The *rpm1-3* (*rpm1*) (Grant et al., 1995), *rps2-101C* (*rps2*) (Mackey et al., 2003), *rps2 rin4* (Mackey et al., 2003), and *rps2 rin4 rpm1* (Belkhadir et al., 2004) mutants were crossed to *sober1-3* (Choi et al., 2018; Cunnac et al., 2007) (*sober1*) to generate *sober1 rpm1, sober1 rps2, sober1 rps2 rin4*, and *sober1 rps2 rin4 rpm1*, respectively. Wild-type Ct-1 was crossed to the *rps2 rin4 rpm1* mutant to generate *sober1*_*Ct-1*_ *rps2 rin4*, which was used to generate the transgenic lines expressing *RIN4* variants. All expected mutations were verified using a genotyping PCR (for primers, see Table S2). The *Arabidopsis* plants were grown at 22°C under short-day conditions (11 h light and 13 h dark). *Nicotiana benthamiana* plants were grown at 24°C in long-day conditions (16 h light and 8 h dark).

### Plasmid constructs

The plasmid constructs used in this study is described in Supplemental Experimental Procedures.

### Hypersensitive response and bacterial growth restriction assay

Broad-host-range vector constructs were mobilized into *Pseudomonas* strains by triparental mating using *E. coli* HB101, as described previously (Jayaraman et al., 2017). For the HR assay, *P. fluorescens* Pf0-1 (T3S) (1 × 10^8^ CFU/mL) was infiltrated into leaves of 4–5-week-old *Arabidopsis* plants. HR symptoms were scored and photographed at 1 dpi. For the *in planta* bacterial growth restriction assay, *Pto* DC3000 strains were resuspended in 10 mM MgCl_2_ to 1 × 10^5^ CFU/mL, and infiltrated into leaves of 4–5-week-old *Arabidopsis* plants. At 4 dpi, six technical replicate samples (*n* = 6) of leaf tissue (1 cm^2^) were harvested and ground in sterile 10 mM MgCl_2_, serially diluted, and plated on King’s B plates containing the appropriate antibiotics. Colonies were counted on the plates after two days of incubation at 28°C.

### Genetic analysis of the *Arabidopsis* F_*2*_ population using next-generation whole-genome sequencing

*Arabidopsis thaliana* Ct-1 and Tscha-1 were crossed to generate an F_2_ population for HopZ5 *R* gene mapping. The HopZ5-triggered HR was scored in 140 F_2_ individuals and the HR segregation ratio was assessed. Genomic DNA was extracted from 37 F_2_ plants that did not show a HR using a DNeasy Plant Mini Kit (Qiagen, Germany). This DNA was pooled and sequenced by DNA Link (Republic of Korea) using an Illumina Novaseq6000 (Illumina, USA). Details of bioinformatic analysis are described in Supplemental Experimental Procedures.

### Generation of transgenic plants

Binary constructs were electroporated into *A. tumefaciens* strain AGL1. To generate transgenic lines in the *Arabidopsis* Tscha-1 background, flowering plants were dipped with *Agrobacterium* carrying pICH86966:*RPM1:6×HA* (Clough and Bent, 1998). Transformants were selected on Murashige and Skoog (MS) agar plates containing kanamycin (50 μg/mL). To generate transgenic lines in an *Arabidopsis* Ct-1 *rps2 rin4* background, *Agrobacterium* carrying pEpiGreen:*35S:FLAG:RIN4* (wild-type *RIN4* or the *RIN4*^*T166A*^ mutant) was used for floral dip transformation. Transformants were grown on soil and selected after several spray applications of BASTA® solution (Bayer, Germany). Protein accumulation levels were assessed in T_2_ plants using a western blot with α-FLAG antibodies.

### Protein expression and extraction in *N. benthamiana*

*Agrobacterium tumefaciens* AGL1 cultures were grown in L medium at 28°C and centrifuged at 2000 ×*g* for 4 min before being resuspended in infiltration buffer (10 mM MgCl_2_, 10 mM MES-KOH, pH 5.6). Concentrations of bacterial suspensions were adjusted (OD_600_ = 0.1–0.4) in infiltration buffer with the addition of LaCl_3_ (final concentration 2 mM) to prevent cell death, then used to infiltrate 4–5-week-old *N. benthamiana* leaves (El Kasmi et al., 2017). At 2 dpi, six leaf discs (8 mm diameter) were harvested from the infiltrated leaves and placed in 1.5-mL microtubes, ground to a fine powder, and boiled in 200 µL of 2× Laemmli buffer (Laemmli, 1970). If the samples were subjected to Co-IP, 1 g of the infiltrated leaf tissue was ground in 4 mL of extraction buffer (1/2 tablet of Sigma Complete™ mini protease inhibitor (per 15 mL), 0.2% NP-40, 10 mM DTT, 1% PVPP, 10% glycerol, 25 mM Tris pH 7.5, 1 mM EDTA, 150 mM NaCl) then centrifuged at 5000 ×*g* at 4°C for 15 min. The protein extracts were then filtered through Miracloth (Merck, Germany) and subjected to a Co-IP analysis.

### Time-course acetylation assay in *Arabidopsis*

Three or four fully expanded leaves per plant of *FLAG:RIN4*-expressing transgenic *Arabidopsis* were infiltrated with Pf0-1 (2 × 10^8^ CFU/mL) resuspended in 10 mM MgCl_2_ with 2 mM LaCl_3_ using a blunt-end syringe. At 0, 6, and 12 hpi, 0.5 g of leaf tissue was sampled and used for protein extraction. The FLAG-RIN4 protein was immunoprecipitated with anti-FLAG beads and immunoblotted with α-AcT166 antibody.

### Recombinant protein purification from *E. coli*

Protein expression in *E. coli* Rosetta (DE3) cells carrying constructs encoding FLAG-tagged RIN4, effector, or SOBER1 variants in the pOPIN vector was induced with a treatment of 0.5 mM IPTG at 28°C for 3 h. Bacterial cells were then centrifuged at 5000 ×*g*, after which the pellets were resuspended in Cellytic buffer B (Sigma-Aldrich, USA) following the manufacturer’s instruction. A 1-mL aliquot of cell lysate was incubated with 40 µL of anti-FLAG M2 affinity gel (Sigma-Aldrich) at 4°C for 2 h. Samples were washed three times with the protein elution buffer (20 mM Tris-HCl, 150 mM NaCl, 10% glycerol) and eluted by adding 3× FLAG peptide (Sigma-Aldrich) in 60 µL of the protein elution buffer following the manufacturer’s instruction.

### *In vitro* acetylation and deacetylation assay

The cell lysate of *E. coli* expressing FLAG-tagged RIN4 protein was incubated with 20 µL of anti-FLAG M2 affinity gel (Sigma-Aldrich) at 4°C for 2 h. After washing three times with GTEN buffer (10% glycerol, 25 mM Tris pH 7.5, 1 mM EDTA, 150 mM NaCl), 17 µL of FLAG-tagged effector protein extract was added to the beads with 1 µL of 1 mM acetyl-CoA lithium salt (Sigma-Aldrich) and 2 µL of 1 µM IP_6_ (Sigma-Aldrich). The acetylation mix was incubated at room temperature for 2 h and boiled with 5× Laemmli buffer for immunoblotting. For the deacetylation assay, the acetylation mix was then washed three times with protein elution buffer, after which 20 µL of FLAG-tagged SOBER1 variants protein extracts were added. After a 2-h incubation at room temperature, the samples were boiled with 5× Laemmli buffer for immunoblotting.

### Co-immunoprecipitation and immunoblot analysis

Anti-AcRIN4 antibody was raised in rabbit against an acetylated T166 RIN4 peptide (CGADGYTacHIFN). Crude antisera were affinity-purified and their specificity was tested by Abclon (Republic of Korea). Co-immunoprecipitation was performed as previously described (Prokchorchik et al., 2020). Protein samples were separated using SDS-PAGE and immunoblotted with α-GFP-HRP (Santa Cruz, USA), α-FLAG (Sigma-Aldrich), α-MYC (Cell Signalling, USA), or α-AcRIN4 antibodies. Proteins were visualized using a mixture of SuperSignal West Pico and Femto chemiluminescent substrates (Thermo Fisher Scientific). For the loading control visualization, the acrylamide gel after an SDS-PAGE was stained with Coomassie Brilliant Blue (CBB) for *E. coli-*expressed proteins, or the PVDF membranes were stained with Ponceau S after immunoblotting for *in planta-*expressed proteins.

### Mass spectrometry analysis

The acetylation mix samples of bead-bound FLAG-RIN4 and HopZ5-FLAG protein variants were separated using SDS-PAGE and CBB staining. The FLAG-RIN4 protein bands were excised and sent for mass spectrometry analysis (NICEM, Republic of Korea). Details of the mass spectrometry analysis are described in Supplemental Experimental Procedures.

### TRV-mediated protein overexpression assay

*Agrobacterium tumefaciens* AGL1 cells harbouring pTRV2 derivatives were grown in L medium at 28°C, harvested, and mixed with pTRV1 at a 1:1 ratio with OD_600_ = 0.5 for each strain, then infiltrated into the leaves of five-week-old *N. benthamiana* plants. At 2 dpi, the *N. benthamiana* leaf samples were collected, snap-frozen, ground, and thawed on ice in a 3× volume of 20 mM sodium phosphate buffer (pH 7.0). The mixture was centrifuged at 16,000 ×3*g* for 5 min, and the supernatant was rubbed on 5–6-week-old *Arabidopsis* leaves with an application of carborundum powder(Aguilar et al., 2016). At 2 dpi, the virus-infected *Arabidopsis* leaf samples were collected for a protein expression analysis. The cell death phenotype was scored and photographed at 7 dpi.

### Primers

The primers used in this study are listed in Table S2.

## Acknowledgements

We thank Dr. Mark Youles, Dr. Mark Benfield, and Dr. Richard Hughes for providing the pOPIN constructs. We also thank Dr. Jay Jayaraman for critically reading this manuscript. This research was supported by Basic Science Research Programs through the National Research Foundation of Korea (NRF) funded by the Ministry of Education (NRF-2019R1I1A1A01060108) and Korean government (MSIT) (NRF-2018R1A5A1023599 and NRF-2019R1A2C2084705).

## Author contributions

S.C., M.P., H.L., R.G., Y.L., and B.C. performed the experiments. S.C., M.P., and K.H.S. designed the experiments. S.C., M.P., M.K., S.K., and K.H.S. analysed the data. S.C., M.P., and K.H.S. wrote the manuscript.

## Supplemental figures

**Figure S1.**
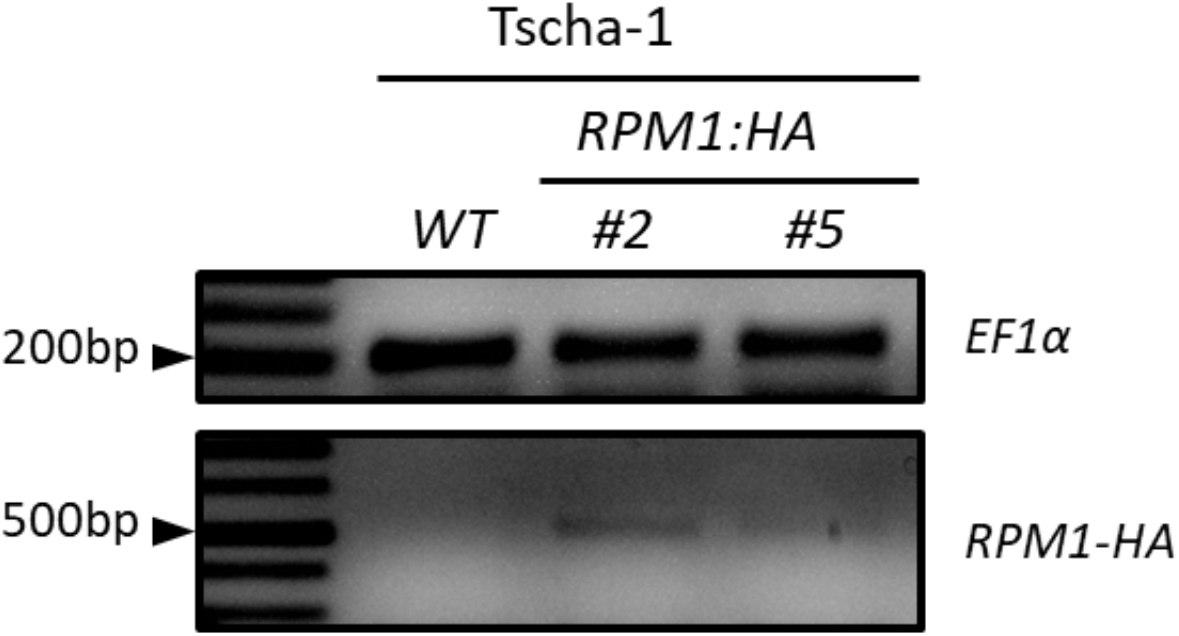
Quantitative RT-PCR for transgene expression analysis in transgenic Arabidopsis Tscha-1 T_2_ lines carrying *RPM1:HA.* *RPM1:HA* transgenic *Arabidopsis* Tscha-1 T_2_ lines used in Fig. 1b and c samples were collected for RNA extraction and cDNA synthesis. *EF1α* and *RPM1:HA* specific primers were used for semi qRT-PCR.

**Figure S2.**
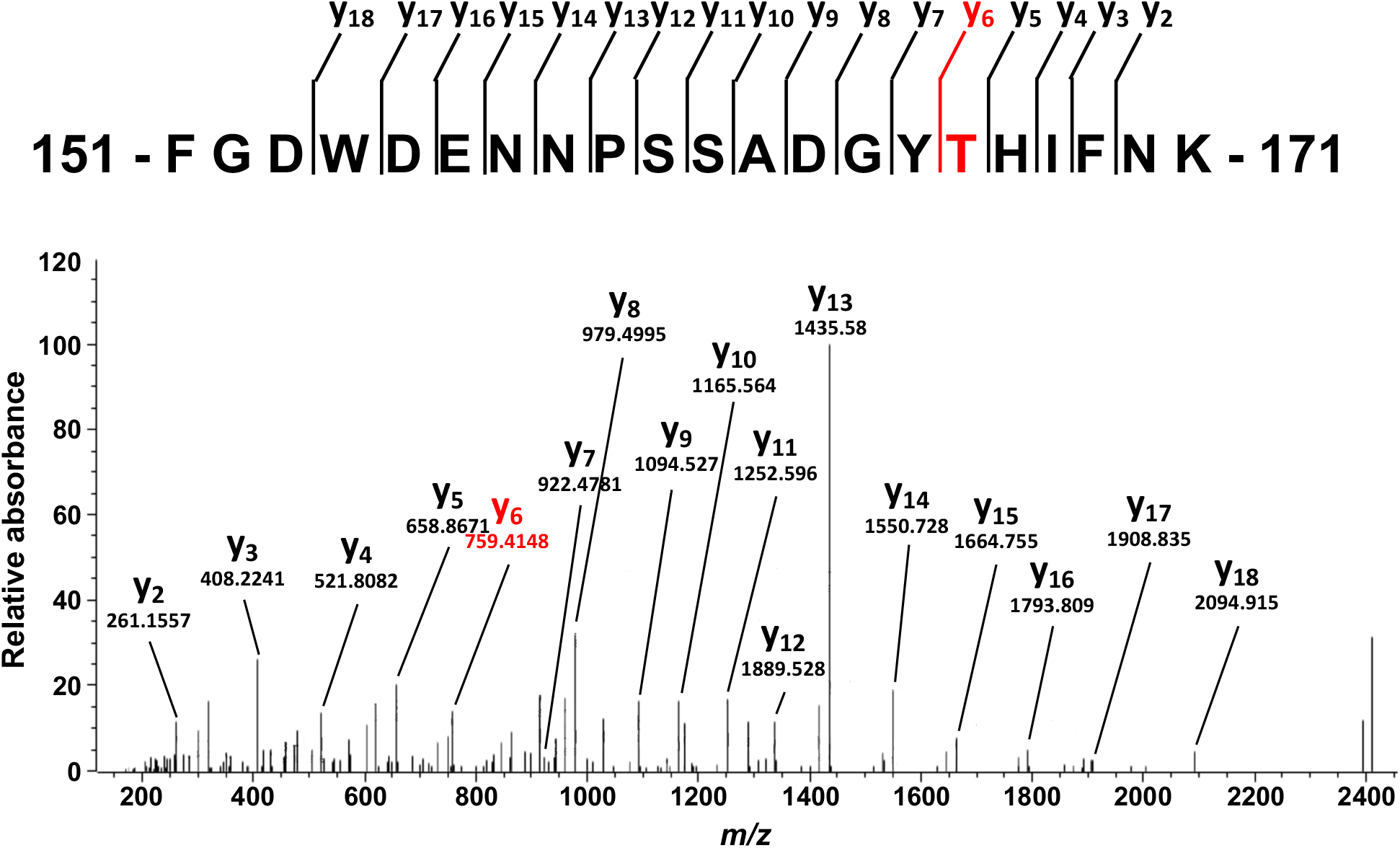
T166 is not acetylated by HopZ5^C218A^ mutant. MS/MS spectra of acetylated RIN4 fragment peptide (FGDWDENNPSSADGYTHIFNK) after incubation with HopZ5^C218A^-FLAG protein. The y ions are marked in the spectrum and illustrated (*m/z*=1229.02). T166 acetylation was not detected after incubation with the catalytic mutant HopZ5^C218A^.

**Figure S3.**
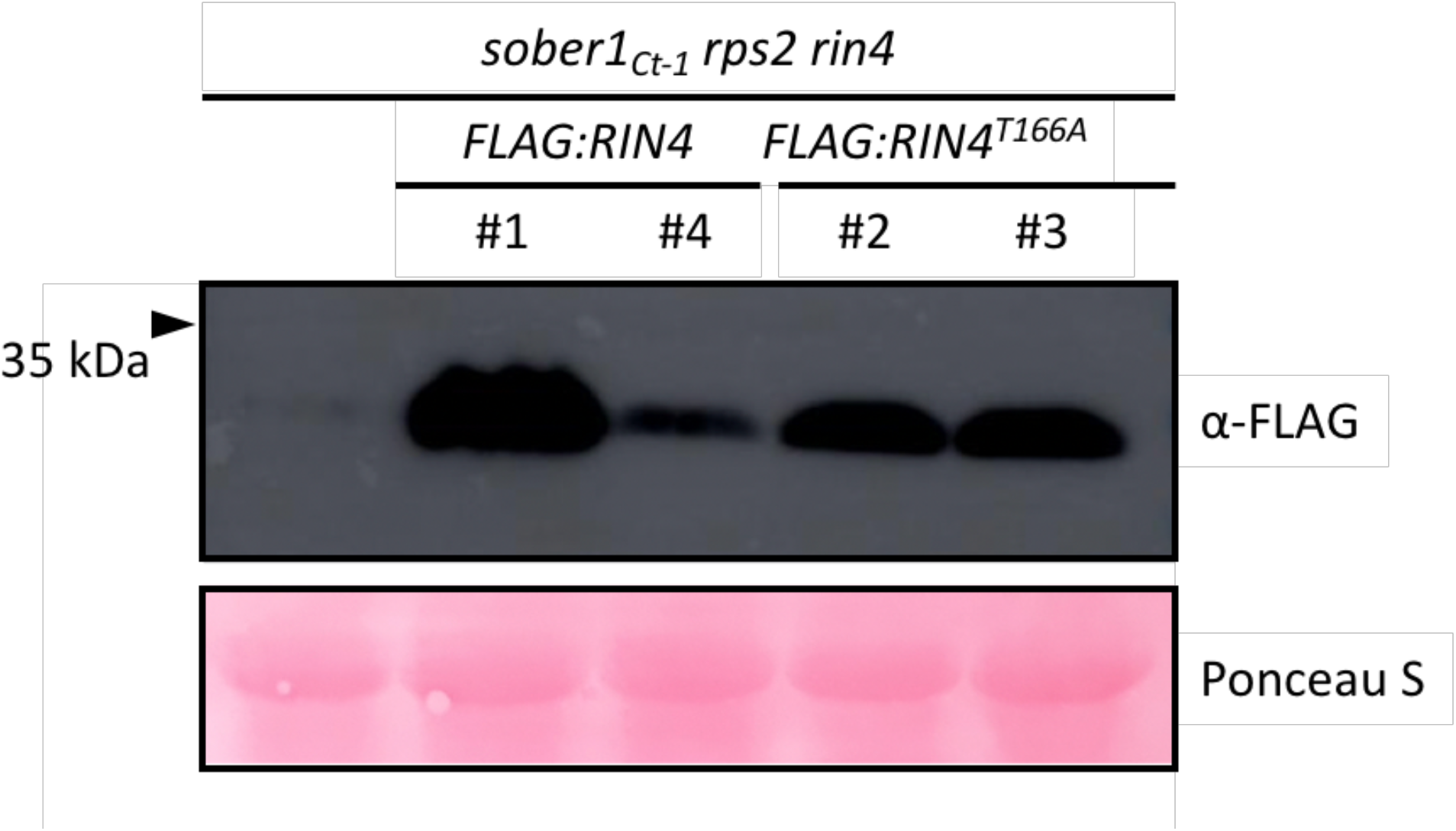
Immunoblot analysis of FLAG-RIN4 and FLAG-RIN4 ^T166A^ proteins in transgenic *Arabidopsis sober1*_*Ct-1*_ *rps2 rin4* lines. Transgenic *Arabidopsis* lines expressing FLAG-RIN4 or FLAG-RIN4^T166A^. Proteins were sampled for total protein extraction and immunoblot analysis. Ponceau S staining of rubisco protein band is a loading control.

**Figure S4.**
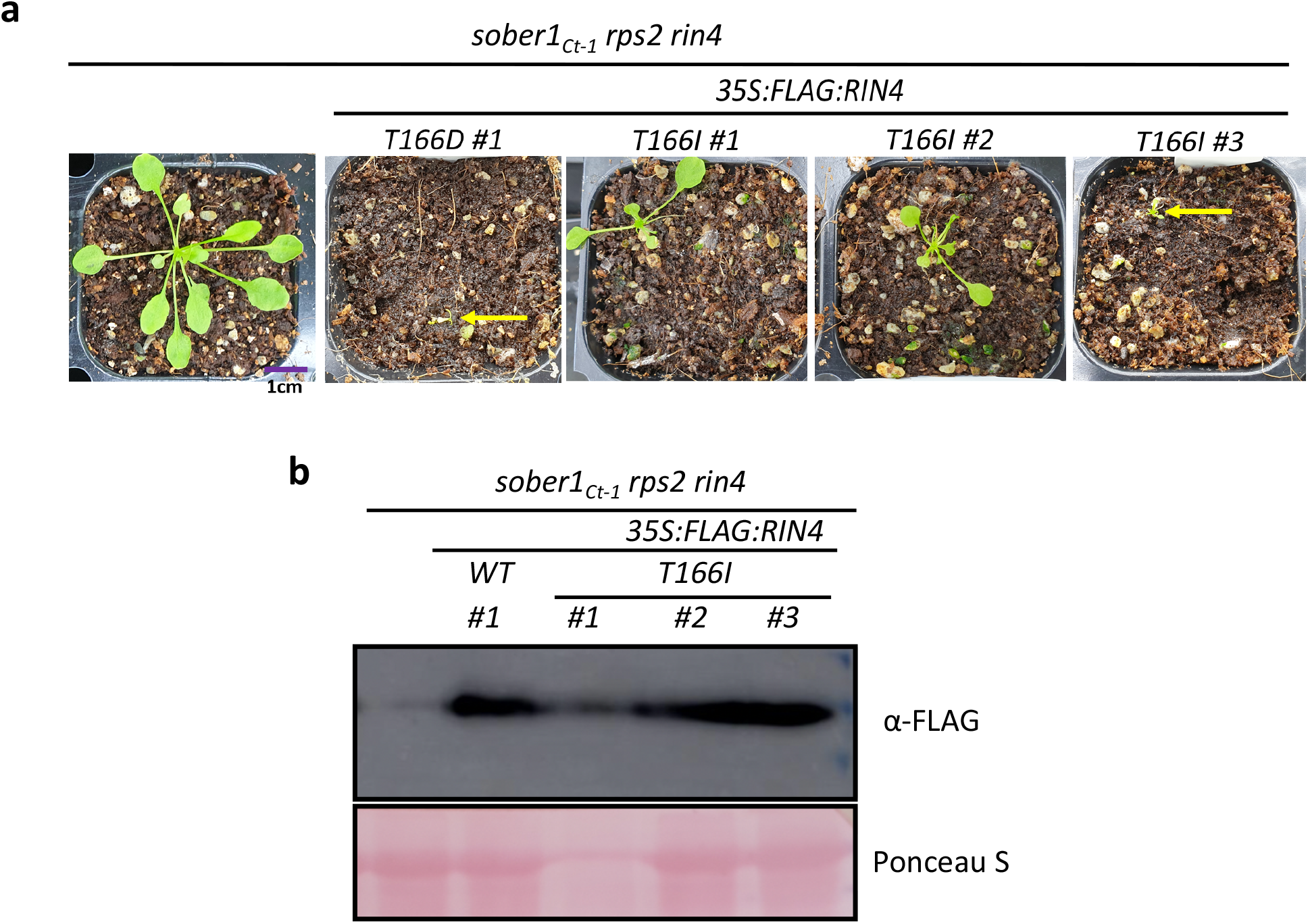
Transgenic expression of RIN4 T166I causes lethality in *Arabidopsis*. **(A)** 4-week old transgenic T_1_*sober1*_*Ct-1*_ *rps2 rin4* lines expressing *FLAG:RIN4 T166D* and *T166I* phenotype was assessed. A purple bar indicates 1 cm. Yellow bar indicates plants. **(B)** Transgene expression in *sober1*_*Ct-1*_ *rps2 rin4 FLAG:RIN4 T166I* T_1_ plants. T_1_ plants were recovered in 28°C for 3 weeks, and the whole plant was sampled for the immunoblot assay. *sober1*_*Ct-1*_ *rps2 rin4* plants and *sober1*_*Ct-1*_ *rps2 rin4 FLAG:RIN4* wildtype T_2_ plant was used as control to compare the transgene expression. Ponceau S staining of rubisco protein band is a loading control.

## Supplemental tables

**Table S2.**
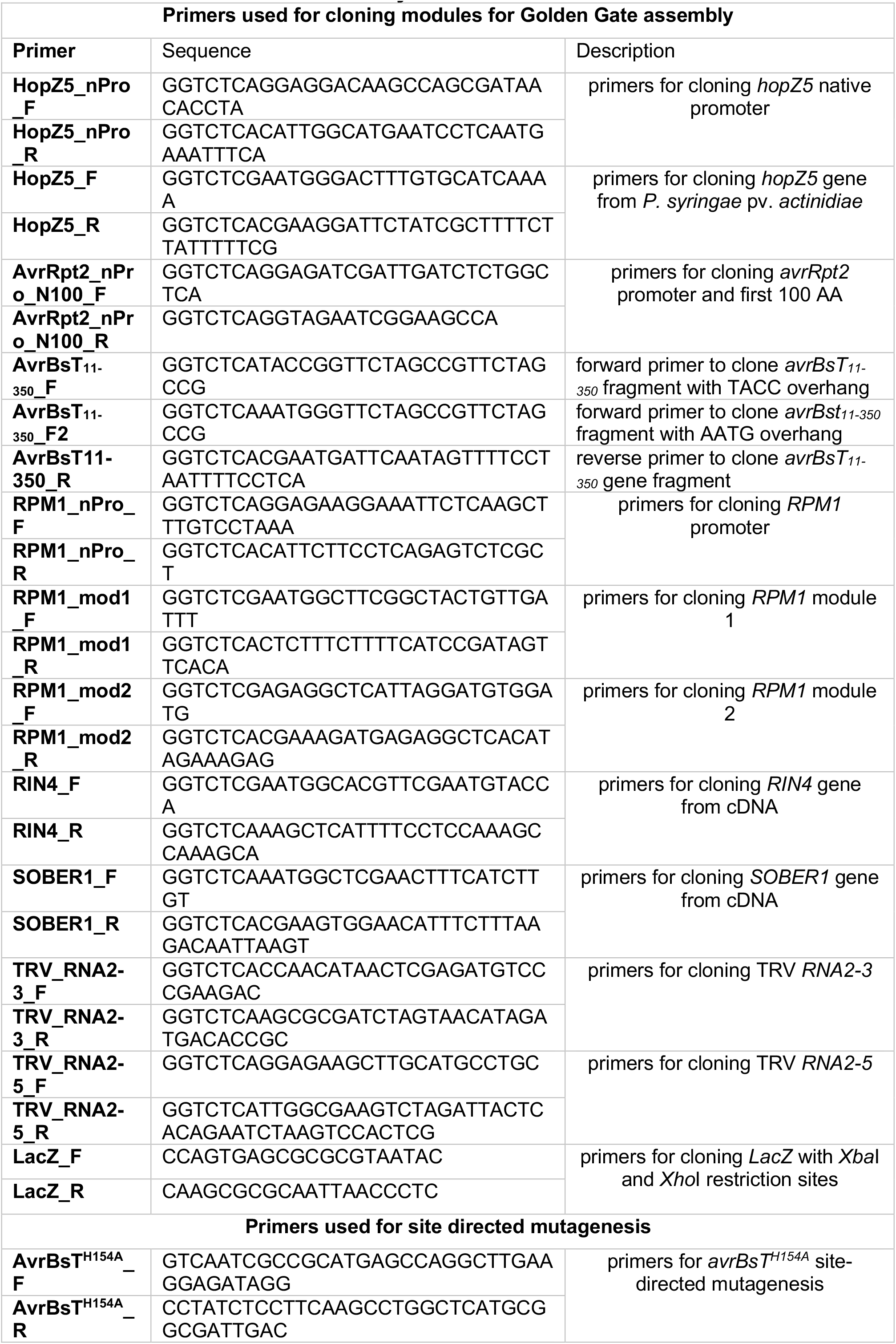
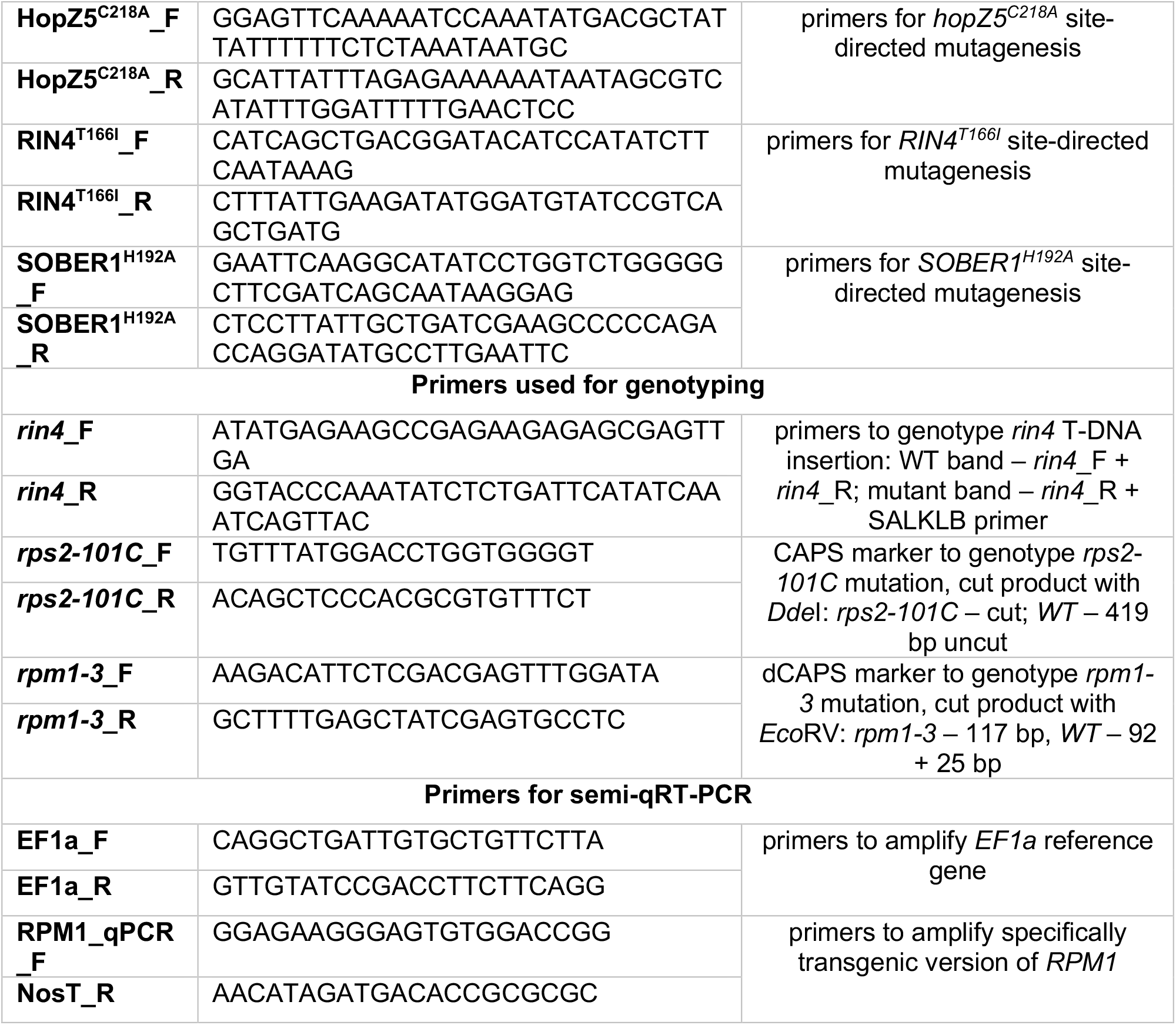
Primers used in this study.

**Table S1.** HopZ5-triggered HR segregation in Ct-1 x Tscha1 F2 mapping population.

| Ct-1 x Tscha-1 F<sub>2</sub><sup>1</sup> | HR<sup>2</sup> | No HR<sup>3</sup> | HR : No HR | $\chi^2$ | p-value |
| --- | --- | --- | --- | --- | --- |
| 140 | 103 | 37 | 3:1 | 0.15 | 0.6985 |

<sup>1</sup> Number of total F2 population tested for HopZ5-triggered HR
<sup>2</sup> Number of F2 population showed HR
<sup>3</sup> Number of F2 population showed no HR

## Supplemental Experimental Procedures

### Plasmid Constructs

The broad-host-range vector constructs carrying *hopZ5* and *hopZ5*^*C218A*^ were generated in the pBBR1MCS-5B backbone under the native 290-bp *hopZ5* promoter region using Golden Gate (Engler et al., 2008) assembly. *avrBsT*^*11–350*^ and *avrBsT*^*11–350 H154A*^ were assembled into the pBBR1MCS-5B backbone under the control of the *avrRpt2* native promoter and fused to the first 100 amino acid residues, as previously described (Choi et al., 2018; Cunnac et al., 2007; Jayaraman et al., 2017). The assemblies of *avrRpt2, avrB, avrPphB*, and *avrRpm1* in the pVSP61 vector were previously described (Ashfield et al., 1995; Innes et al., 1993; Kunkel et al., 1993; Simonich and Innes, 1995). The binary vector constructs of pK7FWG2:*avrRpm1:GFP*, C-terminal YFP-tagged *hopZ5, hopZ5*^*C218A*^, *avrBsT*^*11– 350*^, and *avrBsT*^*11–350 H154A*^ in pICH86988; C-terminally FLAG-tagged *SOBER1* and *SOBER1*^*H192A*^ in pICH86988; and pICH86966:*35S:MYC:RIN4* were previously described (Choi et al., 2018; Jayaraman et al., 2017; Prokchorchik et al., 2020). *RPM1* was cloned under its native promoter (1508 bp upstream of the start codon) into pICH86966 with C-terminal 6×HA epitope tag using Golden Gate assembly (Engler et al., 2008). The DsRed module (Engler et al., 2014) was Golden Gate–assembled with an N-terminal 4×MYC epitope tag under the *35S* CaMV promoter into pICH86966. *hopZ5, hopZ5*^*C218A*^, *avrBsT*, and *avrBsT*^*H154A*^ were Golden Gate–assembled under the *35S* CaMV promoter with C-terminal 3×FLAG in pICH86988. *RIN4* and *RIN4*^*T166A*^ were assembled with N-terminal 3×FLAG tag and under the control of the *35S* CaMV promoter into the Golden Gate–compatible pEpiGreenB5 (Newman et al., 2019). Two versions of kanamycin resistance–containing pOPIN vectors, pOPIN-NT for N-terminal tagging and pOPIN-CT for C-terminal tagging, were used for protein expression in *E. coli* (Berrow et al., 2007). *RIN4* was cloned from *Arabidopsis* Col-0 cDNA and assembled into pOPIN-NT with an N-terminal 3×FLAG tag, and *hopZ5, hopZ5*^*C218A*^, *avrBsT, avrBsT*^*H154A*^, *SOBER1*, and *SOBER1*^*H192A*^ were assembled in pOPIN-CT vectors with a C-terminal 3×FLAG tag. To construct the Golden Gate– compatible pTRV2 vector (pTRV2-66, hereafter), *BsaI-35S:RNA2-5*-*XbaI-BsaI* and *BsaI-XhoI*-*RNA2-3:NOS*-*BsaI* fragments were first PCR-amplified from pYL156 (Liu et al., 2002) and *Sma*I-ligated into pICH41021. Each of the cloned RNA2 fragments was Golden Gate–assembled into pICH86966 to generate pICH886966:*35S:RNA2-5-XbaI-XhoI-RNA2-3*. Subsequently, the *XbaI-lacZ-XhoI* cassette with flanking *Bsa*I restriction sites with specific 4-bp overhangs was amplified and ligated into the pGEM-t-easy TA cloning vector (Thermo Fisher Scientific, USA). pICH886966:*35S:RNA2-5-XbaI-XhoI-RNA2-3* backbone and *XbaI-lacZ-XhoI* cassette were digested with *Xba*I and *Xho*I and ligated to generate the final pTRV2-66 vector. For the virus-mediated protein overexpression, the coat protein (*CP*) gene promoter from Pea early browning virus (PEBV) was synthesized and cloned as a module for Golden Gate assembly (Wroblewski et al., 2018). *RIN4* variants were Golden Gate–assembled under the PEBV *CP* promoter with an N-terminal 3×FLAG tag into pTRV2-66.

### Mass spectrometry analysis

The acetylation mix samples of bead-bound FLAG-RIN4 and HopZ5-FLAG protein variants were separated using SDS-PAGE and CBB staining. The FLAG-RIN4 protein bands were excised and sent for MS analysis (NICEM, Republic of Korea). After a trypsin digestion, the peptides were dissolved again in solvent A (water/ACN, 98:2 v/v; 0.1% formic acid) and separated using reversed-phase chromatography performed on a Dionex UltiMate® 3000 RSLC nano high-performance liquid chromatograph (Thermo Fisher Scientific). To trap the sample, the ultra-high-performance liquid chromatograph was equipped with Acclaim PepMap 100 trap columns (100 μm × 2 cm, nanoViper C18, 5 μm, 100 Å), and the sample was subsequently washed with 98% solvent A for 6 min at a flow rate of 6 μL/min. The sample was continuously separated on an Acclaim PepMap 100 capillary column (75 μm × 15 cm, nanoViper C18, 3 μm, 100 Å) at a flow rate of 300 nL/min. The LC analytical gradient was 2% to 35% solvent B (100% ACN and 0.1% formic acid) over 30 min, then 35% to 95% over 10 min, followed by 90% solvent B for 5 min, and finally 5% solvent B for 15 min. LC–MS/MS was coupled with an electrospray ionization source to the quadrupole-based mass spectrometer Q Exactive™ Orbitrap High-Resolution Mass Spectrometer (Thermo Fisher Scientific). The resulting peptides were electro-sprayed through a coated silica-emitted tip (Scientific Instrument Service, USA) at an ion spray voltage of 2000 eV. The MS spectra were acquired at a resolution of 70,000 (200 *m/z*) in a mass range of 350–1800 *m/z*. The automatic gain control target value was 3 × 10^6^ and the isolation window for MS/MS was 1.2 *m/z*. The eluted samples were used for MS/MS events (resolution of 70,000), measured in a data-dependent mode for the 15 most abundant peaks, in the high-mass-accuracy Orbitrap (Thermo Fisher Scientific) after ion activation/dissociation with higher energy C-trap dissociation at 32 collision energy in a 100–1800 *m/z* mass range. The maximum ion injection times for the survey scan and MS/MS scan were 30 ms and 120 ms, respectively. The acquired MS/MS data were searched with the *Arabidopsis* RIN4 protein sequence and analysed for the presence of lysine, threonine, and serine acetylation using MaxQuant software (Tyanova et al., 2016). Trypsin was specified as a protease, and a maximum of two-missed cleavages were allowed. The carbamidomethylation of cysteine residues was set as a fixed modification, while the acetylation of lysine, serine, threonine, and histidine were set as variable modifications.

### Bioinformatic analysis of next-generation whole genome sequencing

The resulting paired-end Illumina reads with a ∼70× genome coverage were quality-trimmed using fastp (Chen et al., 2018) with default parameters. The reads were then assembled with the *Arabidopsis* Col-0 reference genome using BWA-mem (Li and Durbin, 2009), and the polymorphisms were called using a combination of GATK and Picard components (Van der Auwera et al., 2013). The resulting VCF files detailing the polymorphisms were converted into a SHOREmap-compatible format using the SHOREmap convert command, and the allele frequencies were calculated and analysed using the SHOREmap outcross command with the parameters: window-size: 600,000, window step: 15000, marker min score: 20, marker min coverage: 30 (Schneeberger et al., 2009).

### RNA extraction and semi-quantitative RT-PCR

For RNA extraction 6 leaf discs of 8 mm diameter each were sampled from *Arabidopsis* transgenic lines and snap frozen. Total RNA was purified using Trizol (Sigma-Aldrich, USA) reagent according to the manufacturer’s guidelines. After DNAse treatment, cDNA was generated from 1 μg of extracted RNA using Toyobo ReverTra Ace (Toyobo, Japan) kit according to manufacturer’s manual. *RPM1* transgene specific fragment was amplified and EF1*α* amplification was used as a reference to ensure comparable amount of cDNA in the PCR reaction.

### Immunoblot analysis

For protein sampling whole *Arabidopsis* T_1_ plants (for RIN4^T166D^ and RIN4^T166I^ expressing lines) equalised by weight, or 6 8 mm leaf discs from T_2_ plants (for transgenic lines expressing RIN4 and RIN4^T166A^) were frozen in liquid nitrogen and ground with consequent addition of laemmli buffer. After that protein samples were boiled for 10 min at 96°C and subjected to SDS-PAGE. Proteins were transferred to PVDF membrane, then were treated with anti-FLAG antibodies and visualised using a mixture of SuperSignal West Pico and Femto chemiluminescent substrates (Thermo Fisher Scientific, USA).

